# A novel magnetic bead-based extraction method for the isolation of antimicrobial resistance genes with a case study in river water in Malawi

**DOI:** 10.1101/2021.04.23.439981

**Authors:** Rachel L. Byrne, Derek Cocker, Ghaith Alyayyoussi, M. Mphasa, Mary Charles, Tamandani Mandula, Christopher T. Williams, Jonathan Rigby, Jack Hearn, Nicholas Feasey, Emily R. Adams, Thomas Edwards

**Affiliations:** Centre for Drugs and Diagnostics, Liverpool School of Tropical Medicine. Liverpool, UK; Malawi Liverpool Wellcome Trust. Blantyre, Malawi; Clinical Sciences, Liverpool School of Tropical Medicine. Liverpool, UK; Vector Biology, Liverpool School of Tropical Medicine. Liverpool, UK

**Keywords:** AMR, qPCR, environmental surveillance, river water

## Abstract

**Background:** The environmental is increasingly recognised as an important reservoir of antimicrobial resistance (AMR) genes. Polymerase chain reaction (PCR) and whole genome sequencing (WGS) have great potential in the surveillance of AMR genes. However, molecular methods are dependent upon the isolation of high-quality DNA yields. Currently, there is no consensus for the optimum DNA extraction strategies from complex environmental matrices for downstream molecular applications.

**Methods:** We present a novel magnetic bead-based method for the isolation of antimicrobial resistance genes (ARGs) from river water in Malawi, named MagnaExtract. We present this with analytic limit of detection (LOD) as well as a case study in Southern Malawi. Here we compare the DNA yield and subsequent PCR output from MagnaExtract with commercially available QIAGEN kits and the crude boil and spin method, utilising a high-resolution melt analysis (HRM) PCR panel designed for the detection of third generation cephalosporin and carbapenem resistant genes.

**Results:** Of the 98 water samples evaluated we found the MagnaExtract method to be comparable, and in some instance’s superior to commercially available kits for the isolation of ARGs from river water samples. In addition, we found overnight incubation to promote the recovery of extended spectrum beta-lactamase (ESBL) genes and simultaneous reduction in the detection of carbapenemase genes.

**Conclusion:** The MagnaExtract approach offers a simple, affordable, high yielding extraction method that could be used for the detection of ARGs isolated from river water samples in environmental surveillance campaigns in East Africa.

## BACKGROUND

One of the greatest barriers to addressing antimicrobial resistance (AMR) is access to accurate and reliable diagnostic and surveillance tools. This is highlighted in the World Health Organisation (WHO) global action plan on AMR, which articulates the need for improved diagnostic and surveillance assays in three of the plan’s five strategic objectives (WHO, 2017). One technical approach to AMR surveillance is the development of molecular diagnostics. Nucleic acid amplification approaches such as polymerase chain reaction (PCR) and whole genome sequencing (WGS) can be used to investigate and describe the genotypic profile of bacteria and thus infer their AMR status. PCR and WGS are both, however, critically dependant on the isolation of high-quality DNA (Gupta, 2019; Mantere et al., 2019).

The quality of DNA extracted depends on two main factors: the original sample type and the extraction methods used (Dhaliwal, 2013; Surzycki, 2000). For pure samples such as cultured cells, with sufficient starting material, DNA yield is typically high (Gabor et al., 2003). More complex samples, such as environmental water sources, may contain diverse inhibitors such as salts, DNases and humic compounds that in addition to having a dilution effect, often lead to a vastly reduced DNA yield rendering downstream analysis difficult (Williams et al., 2017).

The environment is increasingly recognised as an important source of AMR genes (Devarajan et al., 2016; Sanderson et al., 2018; Stoll et al., 2012). This is especially true for water sources, with recent studies demonstrating widespread prevalence of ARG in surface water samples (Ng & Gin, 2019; Waseem et al., 2017). A critical question in the epidemiology of AMR is the degree to which there is flux between human, animal, and environmental components. Whilst One Health data are starting to emerge, a large proportion of these studies are set in high-income countries and only report on culturable bacteria in river water (Henriot et al., 2019; Servais & Passerat, 2009; Stoll et al., 2012). However only a small proportion (<0.1%) of aquatic organisms grow on agar media by standard methods (Amann et al., 1995; Stoll et al., 2012). Molecular diagnostics, in particular metagenomic approaches, have the potential to offer an “inclusive” platform to survey the entire diversity of ARGs present in a given sample, as long as adequate DNA can be extracted from these complex matrices. Thus, representing a major advantage over culture-based methods.

Once environmental water has been collected, the next key question is how to process such samples prior to DNA extraction. Firstly, the sample must be concentrated, and there is wide acceptance that water samples should be filtered prior to extraction (Deiner et al., 2015; Eichmiller et al., 2016; Hinlo et al., 2017; Piggott, 2016). There is less agreement about the use of an overnight incubation step in enrichment broth within certain settings despite its regular use in microbiological procedures (da Silva et al., 2012). In the context of AMR surveillance there are pros and cons to this approach; whilst target organisms (i.e. *Escherichia coli*) are effectively amplified, important ARGs on mobile genetic elements (i.e. plasmids) may be lost during this culture step due to a suspected fitness cost (Huang et al., 2013). Furthermore, generalist species like *E*.coli may outcompete niche-adapted pathogens such as *Salmonella* Typhi (Rigby et al., 2021).

Commercially available kits are typically used for DNA extraction from environmental samples, as they offer standardised sets of reagents and are safer than phenol-chloroform-isoamyl alcohol (PCI) extraction methods (Hinlo et al., 2017). Adaptations of manufacturer’s instructions are often reported (Barta et al., 2017; Renshaw et al., 2015), but novel methods are rarely incorporated in high-throughput studies (Oberacker et al., 2019). This has led to inconsistent application amongst environmental researchers and often the kit used is determined by cost, accessibility of materials or personal preference (Hinlo et al., 2017).

Crude DNA extraction methods have been developed such as the boilate technique (boil and spin). The boilate method was established as a low cost and simple process to isolate bacterial DNA (Dashti et al., 2009) from cultured cells in the absence of any chemical reagents or DNA concentration steps. Boilate requires only a heat block for cell lysis and a microcentrifuge to pellet the DNA and remove cellular debris, apparatus available in most settings. However, to our knowledge, its use for complex river water samples has not yet been reported, likely due to inhibitors remaining in the sample.

In recent years, interest in the use of magnetic nanoparticles for DNA purification has increased (Oberacker et al., 2019). Magnetic beads can be coated with a DNA loading antibody or a functional group that specifically interacts with DNA. After binding the DNA, beads are separated from other contaminating cellular components and then purified by ethanol washing. Their utility has, to date, been limited by the high cost of commercially available beads and the lack of open-source methodologies for laboratory developed beads (Oberacker *et al*., 2019). In addition, most available protocols require chemical reagents for lysis and precipitation that can be inaccessible in resource-limited settings.

Here we present an affordable, novel magnetic bead-based extraction method for the isolation of bacterial DNA, and demonstrate its effectiveness using Malawian river water samples, comparing to two commercially available QIAGEN kits and a crude boilate method. Concurrently, we highlight the potential impact of overnight incubation on the recovery of AMR genes from direct and 18-24 hour incubated samples.

## METHODS

### The MagnaExtract method

MagnaExtract utilises sera-mag SpeedBeads (MERCK, Germany) magnetic beads that have been diluted and optimised to be more cost effective than neat, see supplementary material 1 (Fouet et al., 2017; Rohland & Reich, 2012). During development of this protocol numerous factors were considered and tested, outlined in Box 1. Each variable was tested in triplicate using an *E. coli* isolate incubated overnight in buffered peptone water (BPW), a non-selective enrichment broth and evaluated by comparison of cycle threshold (Ct) values using high resolution melt (HRM) PCR for the detection of *bla*_CTXM-1_, and *bla*_SHV_ ARGs, as described by (Edwards et al., 2020). The final method for MagnaExtract is shown in box 2, incorporating the optimised strategy.

#### Box 1

**The optimisation strategy of the MagnaExtract method**. Steps listed in red are variables changed during optimisation. Each variable was conducted in triplicate.

1. Take 200µl of extractant.
2. Heat to 95°C for 10 minutes.
3. Vortex for 15 seconds
4. Centrifuge at 8000RPM for 5 minutes.
5. Transfer 200µl of supernatant to clean 1.5ml Eppendorf
  a. Dilutions made of supernatant to 1:10,1:50 to a final volume of 200µl.
  b. Proteinase K usage
6. Add equal parts magnetic beads.
7. Vortex for 10 seconds
8. Incubate at room temperature on a hula mixer for 5 minutes.
9. Spin down the sample, pellet the beads on the magnetic rack and discard supernatant.
10. Wash with 500µl of freshly made 70% ethanol.
11. Spin down the sample, pellet the beads on the magnetic rack and discard supernatant.
12. Wash with 200µl of freshly made 70% ethanol.
13. Spin down the sample, pellet the beads on the magnetic rack and discard supernatant.
  a. Repeat of step 11 and 12.
14. Air dry for 30 seconds.
15. Remove tube from rack
16. Elute in distilled water
  a. Eluted with 25µl, 30µl, 50µl or 100µl
17. Incubate at room temperature for 2 minutes.
18. Pellet the beads and transfer the supernatant to a clean 1.5ml Eppendorf.

#### Box 2

**The MagnaExtract method**

1. Take 200µl of overnight incubated BPW and isolate solution.
2. Heat to 95°C for 10 minutes.
3. Vortex for 15 seconds
4. Centrifuge at 8000RPM for 5 minutes.
5. Transfer 200µl of supernatant to clean 1.5ml Eppendorf
6. Add 200µl magnetic beads.
7. Vortex for 10 seconds
8. Incubate at room temperature on a hula mixer for 5 minutes.
9. Spin down the sample, pellet the beads on the magnetic rack and discard supernatant.
10. Wash with 500µl of freshly made 70% ethanol.
11. Spin down the sample, pellet the beads on the magnetic rack and discard supernatant.
12. Wash with 200µl of freshly made 70% ethanol.
13. Spin down the sample, pellet the beads on the magnetic rack and discard supernatant.
14. Air dry for 30 seconds.
15. Remove tube from rack
16. Elute in 30µl distilled water
17. Incubate at room temperature for 2 minutes.
18. Pellet the beads and transfer the supernatant to a clean 1.5ml Eppendorf.

#### Analytical limit of detection (LOD)

An *E*.*coli* isolate (ECAB6140/Malawi) was cultured overnight on LB agar at 37°C ±1. One colony was selected and used to inoculate 10ml of buffered peptone water and again incubated overnight at 37°C ±1. A stock solution was then quantified to an approximate concentration of 8 × 10^7^ CFU/ml (OD_600_ 0.1) using a spectrophotometer. A serial dilution series of *E*.*coli* ranging from 10^7^ to 10^0^ CFU/ml was established and 200µl of each dilution was extracted using the MagnaExtract method compared to the DNeasy™ (QIAGEN). Each extraction was performed in triplicate and underwent high resolution melt (HRM) PCR. Each reaction of the HRM assay included 6.25µl of Type-IT 2 x HRM buffer (QIAGEN, Germany), primers specific for the *uidA E*.*coli* housekeeping gene and molecular grade water was added to make a final volume of 12.5µl, including 2.5µl of sample DNA. Reactions were thermally cycled in a RGQ 6000 (QIAGEN), using the profile outlined by Edwards et al. (2018). All analysis was performed in the RGQ software. The LOD was determined by the lowest concentration for which all three extraction replicates amplified.

The Miles, Misra, Irwin method (Miles et al., 1938) was then used to quantify the exact concentration of each dilution by inoculating LB agar plates with 3 × 10µl of each dilution and incubating overnight at 37°C ±1. The 1:100,000 dilution was then used to extrapolate the initial concentration of bacteria in the stock solution to be 5.4 × 10^8^CFU/ml.

### Case study: Detecting ARGs from Malawian river water samples

#### Setting

As part of an ongoing AMR surveillance project, Drivers of Resistance in Uganda and Malawi (DRUM) households are randomly selected based on their geographical location within regions of Southern Malawi (Drum, 2020). Household members are asked to identify their source of river water and sample sites are selected based on their ease of access. Ethical approval for this study was obtained from the University of Malawi College of Medicine Research Ethics Committee (COMREC: P.11/18/2541) and Liverpool School of Tropical Medicine Research and Ethics Committee (LSTM REC: 18-090)

#### Sample collection and processing

Samples were collected in sterile 500ml plastic containers and stored in ice chests, then transported within two hours of collection to our laboratory and stored at 4°C for a maximum of 24 hours prior to processing. All samples were then concentrated using a pump water filtration system of optimum flow rate 3.8-4.0 L/min and passed through VWS Supor® PES membrane filters of aperture 0.45µm (PALL, USA). The filter paper was then cut in two: half was available for immediate DNA extraction and the other incubated overnight in 15ml of buffered peptone water (BPW) at 37°C±1.

#### DNA extraction

Samples were extracted using four different methods: two commercially available kits, the PowerWater and the DNeasy™ blood and tissue kit (Both QIAGEN, Germany); the boil and spin (boilate); and the MagnaExtract method.

DNA was directly extracted from one half of the filter paper using the PowerWater kit following the manufacturer’s instructions, to control for the impact of overnight incubation on ARG recovery. The remaining half was incubated in BPW (Oxoid Limited, UK), after 24 hours 200µl of the incubated BPW was then extracted using the DNeasy™ blood and tissue kit with an additional pre-treatment for Gram-negative bacteria outlined in the manufacturer’s instructions, 200µl using the boilate method and 200µl using MagnaExtract. A 10µl volume of incubated BPW was also used to inoculate a Chromagar™ ESBL plate (CHROMagar, France) and incubated overnight at 37°C±1. A plate sweep was performed by an experienced microbiologist (MM) in order to include all morphologically distinct colonies present on the plate. This was then suspended in 200µl of distilled water, and DNA isolated using the boilate method. For the boilate method the sample was heated to 95°C for 10 minutes, vortexed and centrifuged at 8000RPM for 5 minutes. The supernatant was then retained for downstream application, and the pellet discarded. This was then compared to the MagnaExtract method, outlined in box 2.

#### High resolution melt (HRM) analysis for the presence of antimicrobial resistance genes (ARGs)

Primers for ESBL (*bla*_CTXM-1_, *bla*_CTXM-9_ and *bla*_SHV_) and Carbapenamase (*bla*_IMP_, *bla*_KPC_, *bla*_NDM_, *bla*_OXA-48_ and *bla*_VIM_) genes were taken from a previously published assay (T. Edwards et al., 2020). The threshold value for cycle threshold (Ct) was set at 0.078 dF/dT and retained for all experiments. The presence of carbapenemase genes, was confirmed by an in-house probe-based qPCR assay.

#### Quantitative analysis

The agreement between extraction methods was calculated by comparing the new method (MagnaExtract) to all other methods, for the detection of ARGs within any given sample. For example, if one or more of the extraction methods resulted in the detection of an ARG the sample was deemed to be positive and the MagnaExtract method result was compared to this.

The cost was calculated on a per sample basis to be inclusive of all laboratory consumables and the cost of commercial kits. Electricity and laboratory staffing costs was not included but should be considered.

Data handling, analysis and statistical comparisons were all performed using R (3.5.5) (R, 2020) Statistical analyses for DNA yield were performed using Kruskal-Wallis non-parametric test with Dunn’s post-hoc test to identify differences in yields using each of the five extraction methods. DNA yield was calculated using the Qubit™ 2.0 fluorometer (Thermo Fisher Scientific, USA).

## RESULTS

### Analytical LOD

The analytical LOD in spiked samples indicated the LOD for both MagnaExtract and DNeasy™ (QIAGEN) was 4.0×10^4^ CFU/200µl. The cycle threshold value for each extraction replicate is shown in Table 1. MagnaExtract has a consistently lower Ct value with smaller distributions compared to DNeasy™ (QIAGEN).

**Table 1.** The cycle threshold value of uidA detection in extraction replicates utilising MagnaExtract and DNeasy™ (QIAGEN) methods.

|  | Stock concentration (cfu/ml) | Concentration of extraction volume (cfu/200ul) | Ct value |  |  | Average Ct |
| --- | --- | --- | --- | --- | --- | --- |
| **MagnaExtract** | **2.00E+08** | **4.00E+07** | **22.06** | **21.14** | **22.61** | **21.94** |
| QIAGEN Dneasy™ |  |  | 20.34 | 21.83 | 21.8 | 21.32 |
| **MagnaExtract** | **2.00E+07** | **4.00E+06** | **25.69** | **26.08** | **25.12** | **25.63** |
| QIAGEN Dneasy™ |  |  | 24.78 | 25.76 | 25.48 | 25.34 |
| **MagnaExtract** | **2.00E+06** | **4.00E+05** | **27.98** | **28.69** | **29.33** | **28.67** |
| QIAGEN Dneasy™ |  |  | 28.81 | 28.74 | 29.81 | 29.12 |
| **MagnaExtract** | **2.00E+05** | **4.00E+04** | **29.98** | **30.43** | **30.3** | **30.24** |
| QIAGEN Dneasy™ |  |  | 31.51 | 31.09 | 31.66 | 31.42 |
| **MagnaExtract** | **2.00E+04** | **4.00E+03** | **30.95** | **30.64** |  | **30.80** |
| QIAGEN Dneasy™ |  |  |  | 34.34 |  | 34.34 |

### DNA yield from Malawian river water

The DNA yield was highly variable between the five different extraction methods used. The MagnaExtract method’s overall DNA quantified (mean: 6.78µg/mL, IQR 3.26-13.2) yield was statistically higher than that achieved using DNeasy™, boilate of BPW or boilate of cultured isolate (p < 0.0001), Kruskal-Wallis test, Dunn’s post-hoc test, (n=79) (as shown in figure 2). PowerWater reported similar DNA yield (mean: 4.90µg/mL, IQR 3.56-18.42) with higher variance between samples compared to MagnaExtract highlighted in table 2.

**Figure 1.**
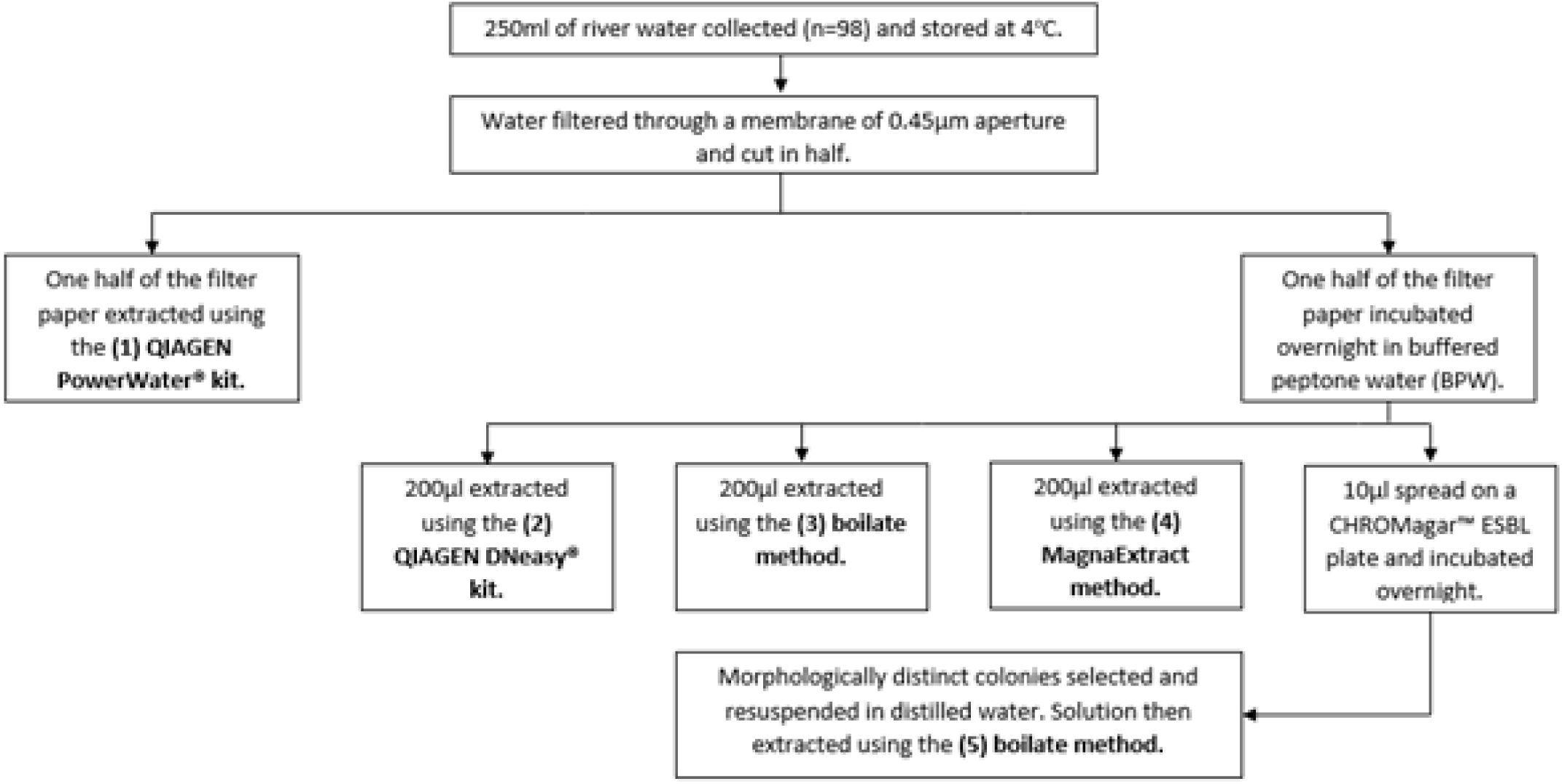
A schematic overview of the methods of DNA extraction utilised. Thus, for all samples, five different DNA samples were obtained, as shown in figure 1.

**Figure 2:**
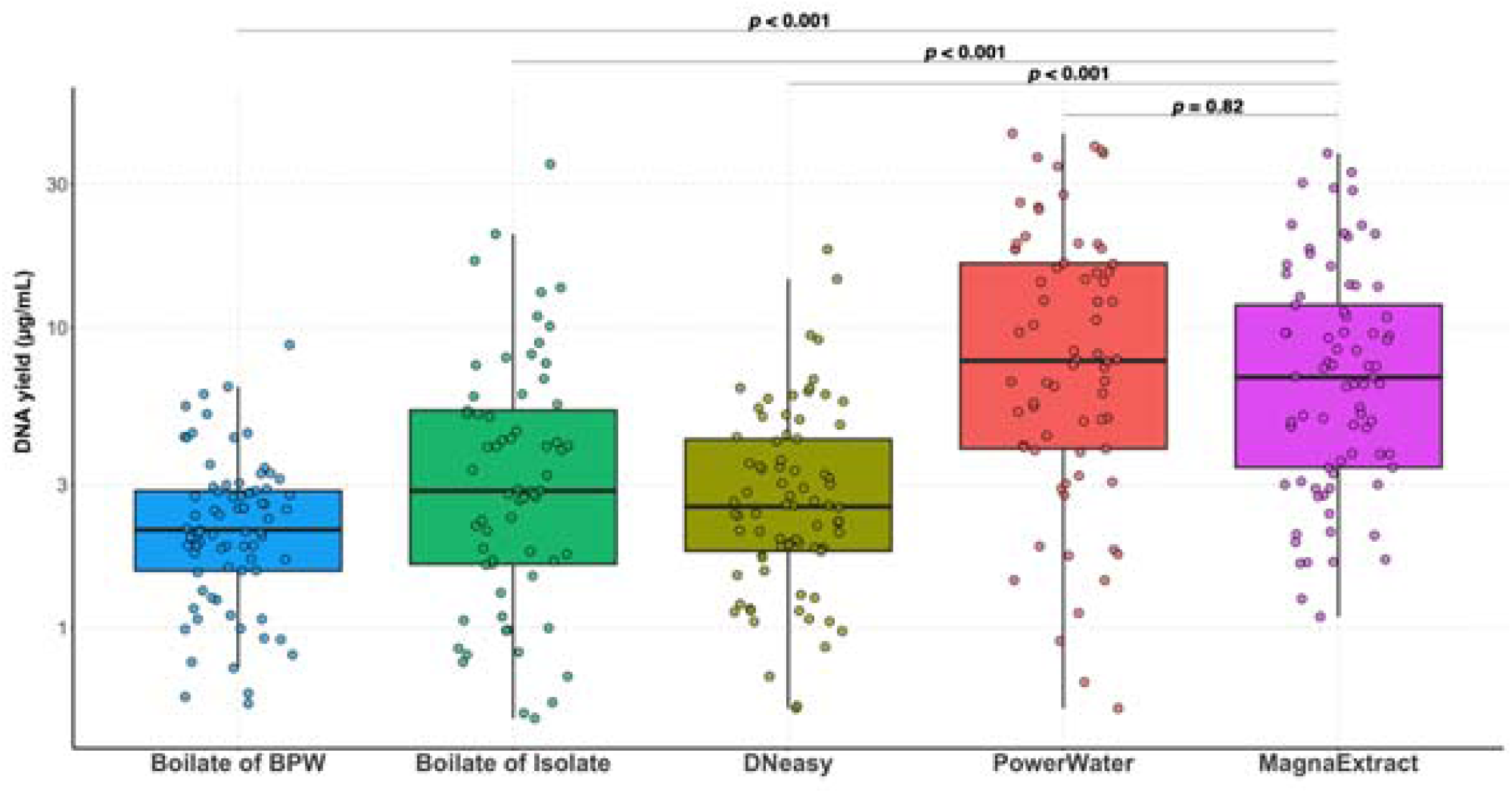
The yield of DNA (µg/mL) from Malawian river water samples (n=79) utilising five different extraction methods. *MagnaExtract is shown to have statistically higher DNA yield than* DNeasy™, *boilate of isolate, boilate and BPW (p < 0.0001) and reported comparable values to PowerWater (p = 0.82), statistical analysis performed using Kruskal-Wallis test with Dunn’s post-hoc test, (n=79)*.

**Table 2:** Comparison of DNA extraction methods for the recovery of bacterial DNA in Malawi river water. Cost of DNA extraction methods calculated on a per sample basis to include cost of commercial kits and all laboratory consumables. Time of extraction method determined from start of extraction to DNA elution. The percentage of positive samples that were correctly identified as positive (agreement), utilising a composite reference standard. Mean DNA yield (as determined by Qubit fluorometer) and standard deviation for all DNA extraction methods for each extraction method.

| Extraction method | Cost (£) | Time (hr) | Agreement (%) | Mean DNA yield |
| --- | --- | --- | --- | --- |
| PowerWater | 8.38 | 1-1.5 | 82 | 13.38 (±14.63) |
| DNeasy™ blood and tissue | 5.38 | 1-1.5<sup>a</sup> | 75 | 3.30 (±2.84) |
| Boilate of BPW | 0.66 | 0.25 <sup>a</sup> | 95 | 2.38 (±1.46) |
| Boilate of isolate | 2.04 | 0.25 <sup>b</sup> | 87 | 4.02 (±5.31) |
| MagnaExtract | 1.43 | 0.5 <sup>a</sup> | 100 | 10.66 (±11.7) |

For time of extraction method from river water collected to DNA elute, <sup>a</sup>24 hours and <sup>b</sup> 48 hours should be added.

### Detection of AMR genes from Malawian river water

Of the 98 river water samples collected, 98.9% (n=97) were positive by PCR by one or more extraction method for ARGs and 92.8% (n=91) positive by plate culture. Only with the MagnaExtract method were all positive samples identified by PCR (table 2) and the lowest sensitivity was reported in commercially available DNeasy™ blood and tissue kit. However, there was little agreement between each method for the positive reporting of ARGs within one sample, as shown in Figure 3.

**Figure 3:**
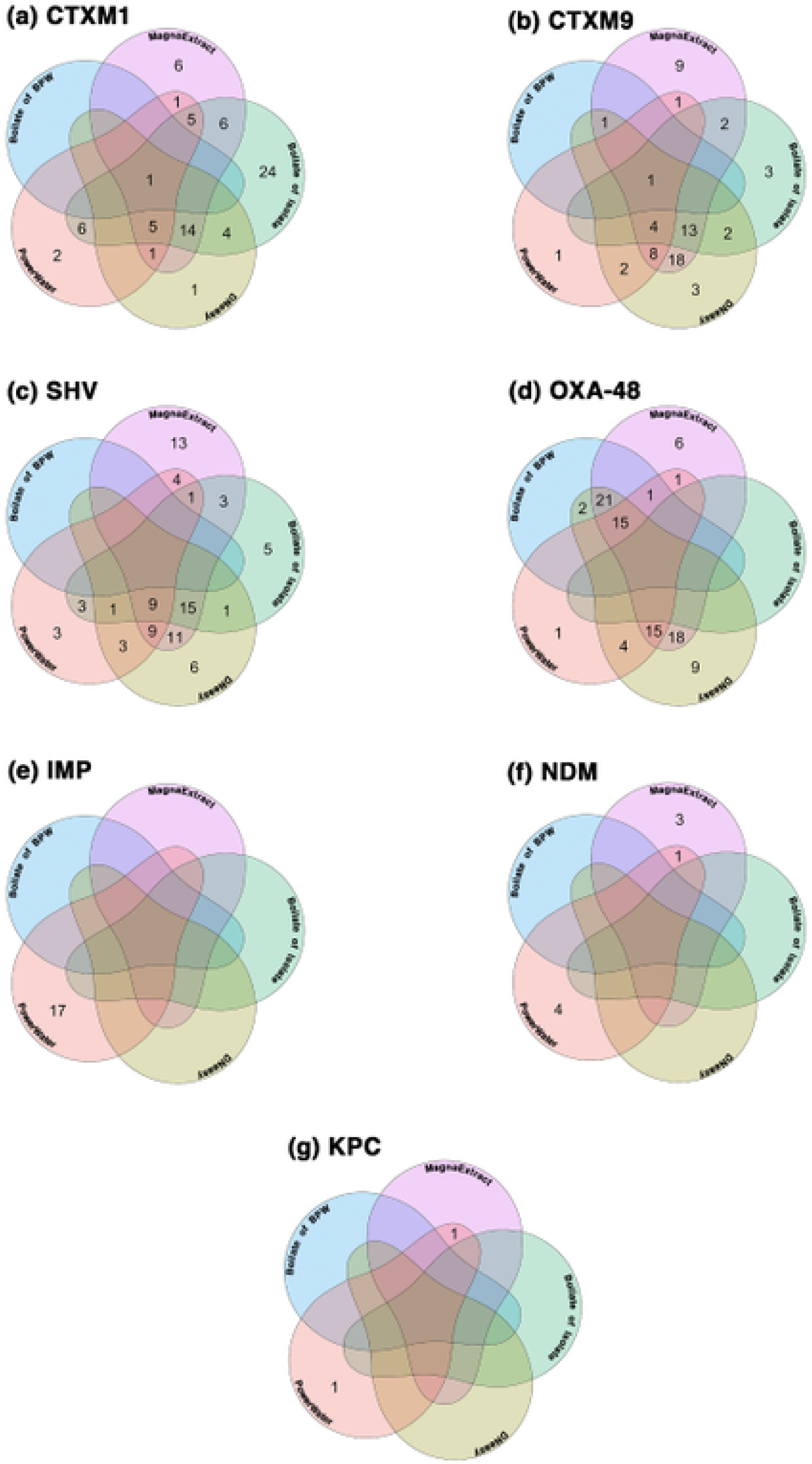
Venn diagrams showing the intersections for the detection of antimicrobial resistance genes (ARGs) extracted from 98 Malawian river water samples using five different methods. *Each Venn is one ARG belonging to either the ESBL class of resistance markers (bla*_*CTXM-1*_, bla_*CTXM-9*_ *and* bla_*SHV*_) or Carbapenamase (bla_*IMP*_, bla_*KPC*_, bla_*NDM*_, bla_*OXA-48*_ *and* bla_*VIM*_)). *Each section of the Venn is a different extraction method used (Commercially available kits PowerWater and* DNeasy™ (QIAGEN, *Germany), crude boilate of BPW, boilate of an isolate grown on ESBL selective media and our novel MagnaExtract magnetic bead-based method*.

OXA-48 (n=258) was the most prevalent ARG followed by the ESBL ARGs (*bla*_CTXM1_, *bla*_CTXM9_ and *bla*_SHV_). The number of ESBL ARGs was greater for methods utilising an overnight enrichment step (DNeasy™, both boilate methods and MagnaExtract). The majority of Carbapenamase ARGs (*bla*_IMP_, *bla*_KPC_ and *bla*_VIM_) were identified by the direct PowerWater kit extraction, and for IMP (n=19) this was the only method by which it was detected. No Carbapenamase ARGs were detected by the boilate of isolate method which involves overnight incubation on ESBL selective growth media, see figure 4.

**Figure 4:**
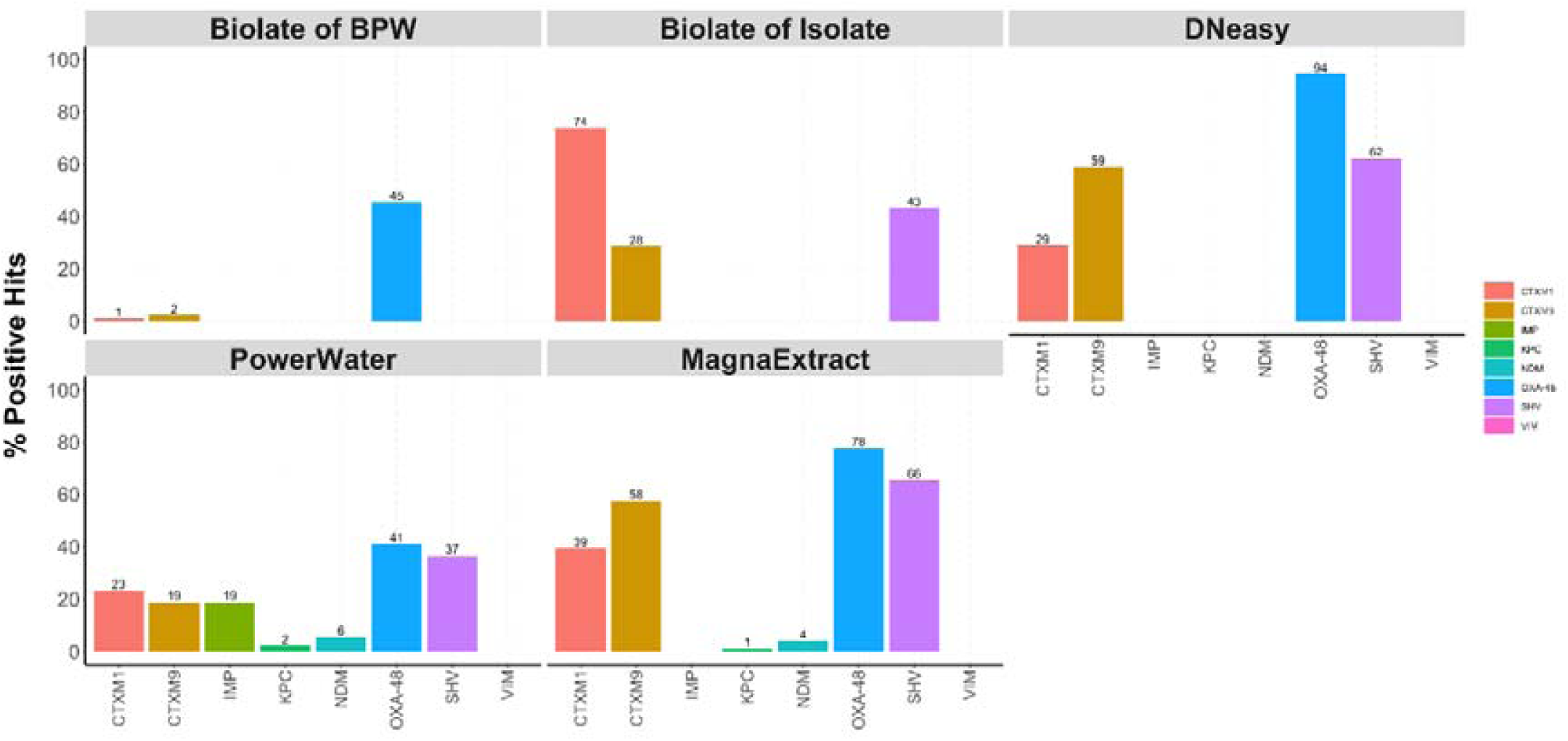
The percentage positive hit of antimicrobial resistance genes (ARGs) isolated from Malawian river water samples using 5 different extraction methods. Each plot depicts the ARGs detectable by each method of interest.

## DISCUSSION

Here we report an inexpensive, high yield, DNA extraction method for the detection of ARGs from river water, a complex environmental matrix employing optimised diluted and buffered magnetic sera-mag SpeedBeads termed ‘MagnaExtract’. This method utilises concentration by filtration followed by overnight incubation in a non-selective growth media (BPW) as is standard practice in many environmental microbiology laboratories globally (Djurhuus et al., 2017). Our novel MagnaExtract method yielded significantly higher amounts of DNA than both commercial and crude methods. The analytical LOD further demonstrated the utility of MagnaExtract in place of the commercial kit as MagnaExtract produced consistently lower Ct values indicative of greater DNA recovery in spiked samples. DNA yield from samples was comparable to direct extraction from the water filter membrane via a commercial kit.

We were able to detect clinically important resistance genes such as *bla*_CTXM-1_, *bla*_CTXM9_ and *bla*_SHV_ from a small volume (250ml) of water using the MagnaExtract protocol. Concurrently, we report the presence of Carbapenemase resistance genes (*bla*_OXA-48_ (n=94), *bla*_IMP_ (n=13) and *bla*_KPC_ (n=3)) in Malawi. It should be noted that, because of the nature of PCR detection, resistance markers cannot be attributed to any specific bacteria species and thus cannot be attributed to clinically relevant bacteria. However, water sources are susceptible to anthropogenic pressures and are often polluted with antibiotics and pathogenic bacteria from human excrement (Sanderson et al., 2018) and can serve as a resistome from which pathogenic bacteria can receive ARGs through horizontal gene transfer (Von Wintersdorff et al., 2016).

We also interrogated the effect of overnight incubation on the recovery of resistance genes. After 24 hours in non-selective growth media, we were unable to detect multiple Carbapenemase resistance genes (*bla*_NDM_, *bla*_IMP_ and *bla*_KPC_). By contrast, a greater number of ESBL resistance genes were detected post incubation than direct from water filter. There has been an increase in data surrounding the relative fitness costs on bacteria that harbour resistance markers in the absence of selection pressure, notably those associated with large mobile plasmids (Cheung et al., 2021; Huang et al., 2013; Melnyk et al., 2015). We hypothesise that the loss here is due to carbapenemases being less stable within bacteria than ESBLs, similar to outcomes observed by Cheung et al., (2021). However, more extensive research is needed to both further our understanding of AMR as well as guide laboratory practice.

The MagnaExtract method of extraction offers an inexpensive and rapid method for the molecular detection of antimicrobial resistance genes from complex river water matrices. It also offers a reliable alternative to expensive commercially available kits with similar, and in some instances, superior DNA yield.

## Acknowledgements

We acknowledge and thank the participants of the Drivers of Resistance in Uganda and Malawi (DRUM). We also acknowledge the valuable contributions of MLW sample collection staff. We thank Professor Charles Wondji for the purchase of magnetic beads

Supplementary material 1: Prepare Homemade sera-mag SpeedBeads

1. Mix sera-mag SpeedBeads and transfer 1ml to a 1.5ml centrifuge tube.
2. Place Speed Beads on magnetic rack until beads are separated.
3. Remove supernatant.
4. Add 1ml TE (pH 7.5-8.0) to beads, remove from magnet, mix by pipetting up and down return to magnet.
5. Remove supernatant.
6. Repeat steps 4 & 5.
7. Add 1ml TE (pH 7.5-8.0) to beads, remove from magnet, mix by pipetting up and down, but DO NOT return to magnet.
8. Add 9g PEG-8000 to a new 50ml conical tube.
9. Add 2.92g NaCl to conical.
10. Add 500ul 1M Tris-HCl to conical.
11. Add 100ul 0.5M EDTA to conical.
12. Fill conical to 49ml using ddH20.
13. Mix conical for 5 minutes until PEG goes into solution.
14. Add 25ul Tween 20 to conical and mix.
15. Add SpeedBeads and TE solution from step 7 to conical and mix.
16. Fill conical to 50ml mark with ddH20.

## Notes

### Competing Interest Statement

The authors have declared no competing interest.

### Summary of Updates

The version of the manuscript has been revised to update the following: The methods is now structured with a clear explanation of the development of the MagnaExtract method. The analytical limit of detection has now been demonstrated.

